# Post-fledging space use and survival in hand-reared versus wild juvenile herring gulls

**DOI:** 10.64898/2026.03.03.709292

**Authors:** Reinoud Allaert, Jolien Van Malderen, Wendt Müller, Eric W.M. Stienen, An Martel, Luc Lens, Frederick Verbruggen

## Abstract

Parental care can shape post-fledging behaviour through provisioning, guidance and social information, yet its absence may alter how young birds establish space use and habitat preferences. We tested the consequences of absent parental care by comparing, hand-reared juvenile herring gulls released without parents with wild, parent-reared conspecifics, focusing on the first two months after fledging. Wild juveniles frequently revisited their natal nest during the first month, whereas hand-reared birds rarely returned to the release site; revisits declined in both groups by the second month but remained more common in wild birds. Wild juveniles used smaller ranges that subsequently expanded, while hand-reared birds began with larger ranges that later contracted, leading to convergence. Contrary to expectation, wild juveniles occurred in areas with higher human population density than hand-reared birds. Habitat use also differed between groups and changed over time. Early on, wild juveniles concentrated activity in anthropogenic and marine habitats, whereas hand-reared birds used rural green habitats more. Later, both groups shifted away from marine areas towards rural green habitats, reducing but not eliminating between-group differences. Short-term survival, did not differ between hand-reared and wild juveniles, indicating that parental care primarily reshaped early space use and habitat choice rather than immediate survival.

## 1 Introduction

Early life is a critical phase in the development of many animals, as the transition from parental dependence to functional independence requires juveniles to rapidly acquire and integrate the behavioural skills needed for survival (Grüebler and Naef-Daenzer, 2010; Marchetti and Price, 1989; Yoda et al., 2004). These skills involve locating and handling food, avoiding predators, and navigating unfamiliar environments. A central process thought to mediate outcomes during early independence is parental care (Jones et al., 2025; Royle et al., 2012). In many taxa, parental contributions before the onset of total independence shape juvenile condition and competence, through provisioning, protection, and behavioural guidance.

This influence is especially well documented in birds, where parental care strongly affects offspring growth and development (Ricklefs, 1968). In many altricial and semi-precocial bird species, parents continue to provide care after young leave the nest or begin moving away from the breeding site. During this stage, juveniles are mobile but still behaviourally inexperienced, and variation in the duration and quality of this extended parental care is commonly associated with higher survival (López-Idiáquez et al., 2018; Szipl et al., 2019; Tarwater et al., 2011). Parents may continue feeding and protecting their young, providing direct energetic support and safety during early independence (Naef-Daenzer and Grüebler, 2016; Tarwater et al., 2011). In addition, they may guide juveniles to profitable foraging sites, shaping early foraging behaviour and space-use patterns via local enhancement or imprinting on foods, places, or cues as competence develops. By repeatedly moving with their young, parents can also constrain early dispersal, promote site fidelity, and bias juveniles towards familiar areas and routes during the first weeks of independent movement (Overveld et al., 2011). For example, adult pied avocets (*Recurvirostra avosetta*) regularly lead broods from nesting sites to the most profitable feeding areas (Lengyel, 2006). By observing parents, juveniles can also learn food-handling skills or the use of tools, as demonstrated in, for example, New Caledonian crows (*Corvus moneduloides*) (Holzhaider et al., 2010).

When extended parental care is reduced or absent, juveniles may have to rely more on trial-and-error learning, which can lead to lower early foraging competence, poorer predator recognition, and fewer social-information pathways (Curio et al., 1978; Marchetti and Price, 1989; Yoda et al., 2004). It may also lead to earlier dispersal and wider ranging, because juveniles are not guided back to profitable or safe sites and must locate resources independently. Reintroduction programmes offer an illustrative case of such conditions. Depending on the approach, young may have received parental care before fledging, but sustained post-fledging support is rarely possible because parents do not accompany juveniles after release, and in headstarting juveniles develop without parental care throughout (Buner et al., 2011; Donaldson et al., 2024). For example, reintroduced bearded vultures (*Gypaetus barbatus*) range more widely than wild conspecifics (Margalida et al., 2013), and reintroduced Bonelli’s eagles (*Aquila fasciata*) disperse earlier and revisit natal areas less frequently than juveniles raised with parents (Egea-Casas et al., 2023). Together, these patterns reinforce the idea that parents shape early space use and the tendency to remain in or return to familiar sites of their chicks during the initial period of independent movement.

Despite these indications, well-powered field comparisons of juveniles developing with or without natural parental care remain scarce (Jones et al., 2025), because parental care is hard to experimentally manipulate, natural variation is confounded with factors such as parental age and condition, brood size and hatch date, and sustained monitoring of early post-fledging movements and space use remains technically challenging in many systems.

The herring gull (*Larus argentatus*) offers a highly suitable model system to overcome these limitations. It is a long-lived larid with extended post-fledging parental care: adults provision chicks at the nest for 5–7 weeks (Risch and Rohwer, 2000) and commonly continue to feed and attend juveniles for several more weeks after they have left the natal area (Burger, 1981; Graves et al., 1991; Holley, 1982). Juvenile behavioural competence develops gradually throughout this period, consistent with the slow life-history strategy of large gulls, which is characterised by delayed maturity and a prolonged developmental window for acquiring foraging and movement skills through practice and social learning (Gamero and Kappeler, 2015; Morel et al., 2024; Slagsvold and Wiebe, 2011). This model system provides two additional key advantages. First, herring gulls are well-suited to experimental hand-rearing, unlike many more sensitive taxa. They can be successfully raised without parental care in large, standardised cohorts. This enables a robust direct comparison between naturally reared juveniles and those reared by humans. Second, their relatively large body size allows GPS deployment at a young age (before they leave the nest area) making it feasible to quantify the effects of parental care deprivation on early movement and space use in sufficiently large sample sizes.

Recent work in the same study area provides a useful baseline for interpreting early-life contrasts in herring gulls and the closely related lesser black-backed gull (*Larus fuscus*). Using long-term ringing and resighting data from >3,400 wild and rehabilitated juveniles, we found subtle behavioural differences during the juvenile stage: rehabilitated gulls were resighted more frequently in intertidal areas and in zones with lower human population density than wild conspecifics. These differences diminished as birds matured and space-use patterns converged and survival rates did not differ significantly between rehabilitated and wild birds across juvenile, immature, and adult stages (Uytterschaut et al., 2025).

However, the rehabilitated juveniles in the previous study had already received substantial early parental care prior to admission to the rehabilitation centre, which could have a major impact on their movement and survival. Moreover, resighting-based analyses are constrained by coarse spatial and temporal resolution, preventing detailed inference about fine-scale movements or the mechanisms through which early-life conditions structure space use. The present study addresses these issues by using high-resolution GPS tracking to compare the first two months after leaving the natal site in naturally reared wild juveniles versus hand-reared juveniles that received no parental care. To make this contrast as clear as possible, we designed the rearing protocol to minimise potential effects of captivity unrelated to the absence of parental care.

Because parental attendance can guide early movements toward profitable or familiar sites and expose young to specific foods, places and cues, we predicted behavioural differences between groups. Specifically, we expected hand-reared juveniles to (1) spend less time near their site of origin, because wild-reared juveniles can continue to receive parental provisioning around the nest and colony area after fledging; (2) range more widely over time, because the absence of parental guidance and social information removes a comparable spatial anchor and may force juveniles to search more broadly to locate suitable foraging and roosting sites; and (3) occur more frequently in areas with higher human population density, reflecting responsiveness of hand-reared individuals to human carers during rearing. We also anticipated differences in (4a) overall habitat use and (4b) fine-scale habitat selection, as the presence or absence of parental guidance can influence how juveniles learn to exploit different environments. However, because these patterns may depend on local conditions and individual experience, we did not specify *a priori* directional expectations. Finally, because parental care can buffer early risks and support the development of foraging competence, we further predicted (5) lower short-term survival in hand-reared juveniles. This study asks whether juveniles reared without parents differ from naturally reared juveniles in origin-site use, ranging behaviour, habitat use and selection, and short-term survival during the first two months after fledging.

## 2 Methods

### Experimental groups

#### Hand-reared individuals

During May and June 2022–2024, herring gull eggs were collected from roof nests along the Belgian coast by the Research Institute for Nature and Forest (INBO) and the ‘gull patrol’ team under licence from the Agentschap voor Natuur en Bos (ANB) as part of nuisance-prevention measures. Eggs were transported to Ghent University’s avian facilities (Lab no. LA1400452) at the Wildlife Rescue Centre (WRC) in Ostend. Upon arrival, they were assigned unique identifiers, and incubated at 37.5°C (45% humidity) until pipping, after which incubation continued at 37.2°C (50% humidity) in a hatchery.

After hatching, the birds were housed in groups of 10 in heated boxes (120 × 60 × 60 cm; ambient 15–25 °C; 40–80 % humidity) under natural photoperiod, with heating plates (30 × 30 cm) until day 5 and fed small pieces of fish, soaked dog pellets, and Akwavit supplement (Kasper Faunafood, The Netherlands) *ad libitum*. From day 5, chicks above 60 grams were moved to outdoor enclosures (500 × 205 × 265 cm) in stable groups of 8–10, with heating plates provided when nighttime temperatures fell below 5 °C or during adverse weather. Feed (dog pellets and fish) was provided four times daily; water remained available *ad libitum*.

Approximately five weeks prior to release, juveniles were moved into a larger and higher flight cage (approx. 180 m2) than the original enclosures, allowing them to practise flight skills and dehabituate them from handling. To reduce effects related to predictable provisioning and human association (Willette et al., 2023), feeding was then reduced to once per day at irregular times and in variable amounts, with access often delayed by freezing fish or pellets in water, and food was provided from behind shade cloth so that the birds could no longer see the animal caretakers.

Once juveniles reached fledging age and demonstrated sustained, controlled flight, they were released at the IJzermonding nature reserve (51°09’ N, 2°43’ E). A non-random subset (150 individuals out of 204) was equipped with a GPS tag (see below), based on a qualitative pre-release assessment by experienced staff that prioritised birds with intact, well-developed flight feathers, robust body condition, and strong flight performance. Individuals were approximately 10 weeks old at release.

#### Wild-reared individuals

In 2023 and 2024, we monitored 175 wild-reared chicks at rooftop sites in Ostend (51°13’ N, 2°56’ E) and at the ground-breeding colony in Zeebrugge (51°20’ N, 3°10’ E). Nests were visited three times per week throughout the breeding season, allowing chick age to be tracked from known hatch dates. By day 30 post-hatching, 149 chicks were still alive. We targeted chicks approaching fledging age, when they would soon become difficult to capture. Chicks were eligible for tagging once they were at least 29 days old. To minimise disturbance and non-independence within broods, we aimed to tag a single chick per nest, selecting the individual judged to be in the best condition at the time of capture based on a qualitative assessment of feather development and body condition. Selected individuals were captured by hand and fitted with a GPS transmitter.

#### GPS tracking

Across the three breeding seasons, 204 juveniles (150 hand-reared; 54 wild) were equipped with solar-charged GPS transmitters (Interrex FLEX II 3G/4G) attached via Teflon ribbon body-harnesses (Thaxter et al., 2014). Devices (including harness) weighed approximately 19 g (< 3 % of the lightest bird’s mass) and were programmed identically for both groups. Location was recorded every 30 min, with increased frequency under optimal solar conditions. Every 30 min the units also collected 30 s of 25 Hz accelerometer data, from which onboard calculations provided overall dynamic body acceleration (ODBA). Six individuals whose tags never transmitted any data after deployment were excluded from further data processing.

Analyses of GPS data were conducted relative to the moment of fledging, defined as the first calendar day on which a bird moved more than 500 m from its *origin site* (i.e., the nest for wild-reared individuals and the release site for hand-reared individuals). Across years, wild-reared juveniles reached this threshold on average on 31 July (range 11 July–17 August), whereas hand-reared juveniles did so on average on 13 August (range 9–26 August). For each individual, we extracted GPS data for the subsequent 60 days. This interval was chosen because preliminary checks showed that most juvenile mortality occurred within this period, and because solar recharge reliably sustained 30-minute GPS fixes during these 60 days, whereas beyond this point declining light levels reduced recharge and led to inconsistent sampling rates. We then split this 60-day window into two consecutive 30-day blocks (days 1 to 30 and 31 to 60 post-fledging) to test whether behaviour changed over time.

#### Software and model diagnostics

All analyses were conducted in R 4.3.2, with package versions tracked via `renv` (Ushey and Wickham, 2024). Dispersion, zero-inflation, and residual diagnostics for count models were assessed with `DHARMa;` additional checks of normality, homoscedasticity, and autocorrelation, as well as overall fit, used the `performance` package (Lüdecke et al., 2021). Post hoc contrasts and estimated marginal means were generated with `emmeans`(Lenth, 2025), with multiplicity adjustments as appropriate.

### Behaviour (Predictions 1-4)

To compare the behaviour of both groups, we included only individuals with a complete 60-day period of continuous, high-resolution GPS data. Continuity was determined by calculating inter-fix intervals for each bird and excluding individuals whose mean interval exceeded 35 min or whose maximum gap between fixes exceeded 24 h. All remaining tracks were then subsampled to a regular 30-min interval to standardise sampling effort across individuals. After filtering, 87 birds were retained for behavioural analyses (65 hand-reared, 22 wild-reared; see Table 1).

**Table 1.** Summary of fate classifications for juvenile herring gulls, indicating the total number of individuals assigned to each category by origin (hand-reared vs. wild-reared).

| Category | Description | Sample size |  |
| --- | --- | --- | --- |
|  |  | Hand-reared | Wild-reared |
| Tag malfunction | Tag failed shortly after deployment; no usable data. | 6 | 0 |
| Confirmed dead | Movement ceased and accelerometer profiles flatlined, or carcass recovered. | 66 | 15 |
| Potentially dead | Tag stopped transmitting within the 60-day window; fate uncertain. | 12 | 2 |
| Alive | Continuous GPS fixes and dynamic accelerometer profiles throughout the 60 days. | 66 | 37 |

Because most foraging and active movement occur during daylight hours (Baert et al., 2022), only GPS fixes that occurred during daytime (from one hour before sunrise to one hour after sunset) were retained for all behavioural analyses. These fixes were identified using the `suncalc` package (Thieurmel and Elmarhraoui, 2022). After filtering, fixes were annotated with habitat (CORINE Land Cover 2018; Copernicus Land Monitoring Service, 2019) and human population density (2020 GHS-POP; European Commission, Joint Research Centre, 2023) rasters, using the `terra` package (Hijmans, 2025).

#### Prediction 1: Time spent at site of origin

To determine time spent at the site of origin, we first calculated distances for each fix to the origin site using the `geosphere` package (Hijmans, 2024). All GPS positions within 500 m of the origin site were grouped into a single “origin site” area. This 500-m threshold was used to aggregate fine-scale movements around the nest or release location (which individuals could always return to).

We then counted all daytime fixes in the origin-site area for each bird in each of the two 30-day blocks after fledging (days 1–30 and 31–60), and fitted a binomial GLMM (logit link) with fixed effects of *group* (hand-reared vs. wild-reared), *month block* (first vs. second 30-day block) and their interaction, and a random intercept for individual, using `glmmTMB` (McGillycuddy et al., 2025). Because GPS analyses began only after individuals first moved >500 m from their origin site, “time spent at the origin site” refers to subsequent revisits to the nest (wild-reared birds) or release site (hand-reared birds) during the 60-day post-fledging period.

#### Prediction 2: Space use

We quantified space-use dynamics by estimating 95% minimum convex polygons (MCPs) in rolling 10-day windows using the `amt` package (Signer et al., 2019), see Figure 1 for an example. For each bird, a 10-day window was slid along its 60-day post-fledging track (i.e., day 1–10, 2–11, …, 51–60), and only windows containing at least 100 GPS fixes were retained for MCP estimation. Within each qualifying window, the 95% MCP was computed via `hr_mcp()`, its area (in hectares) extracted using `hr_area()`, and then log-transformed to normalise variance. We chose MCPs because they delimit the convex hull of observed locations, providing a conservative estimate of the actual area used within each 10-day window rather than extrapolating beyond recorded fixes.

**Figure 1.**
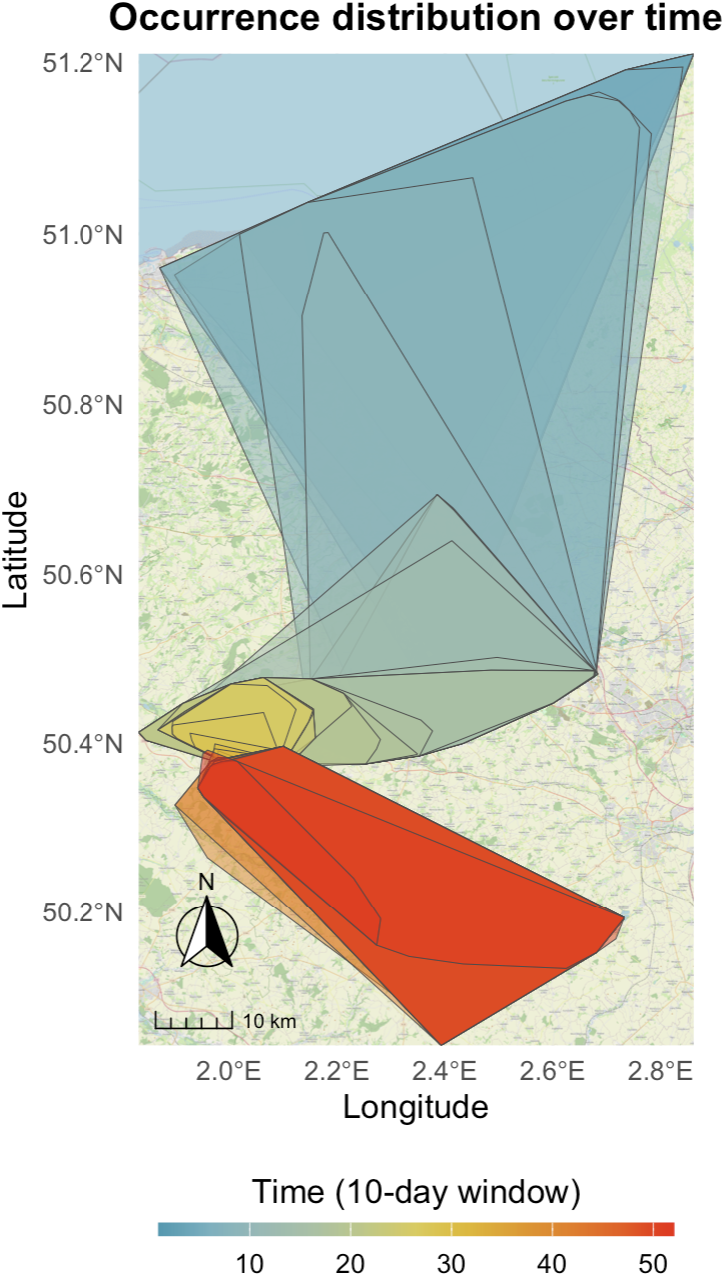
Example of rolling 10-day 95% Minimum Convex Polygons (MCPs) for a juvenile herring gull. Each semi-transparent polygon represents the 95% MCP of all GPS fixes collected within a consecutive 10-day window, with colours indicating the time window.

**Figure 2.**
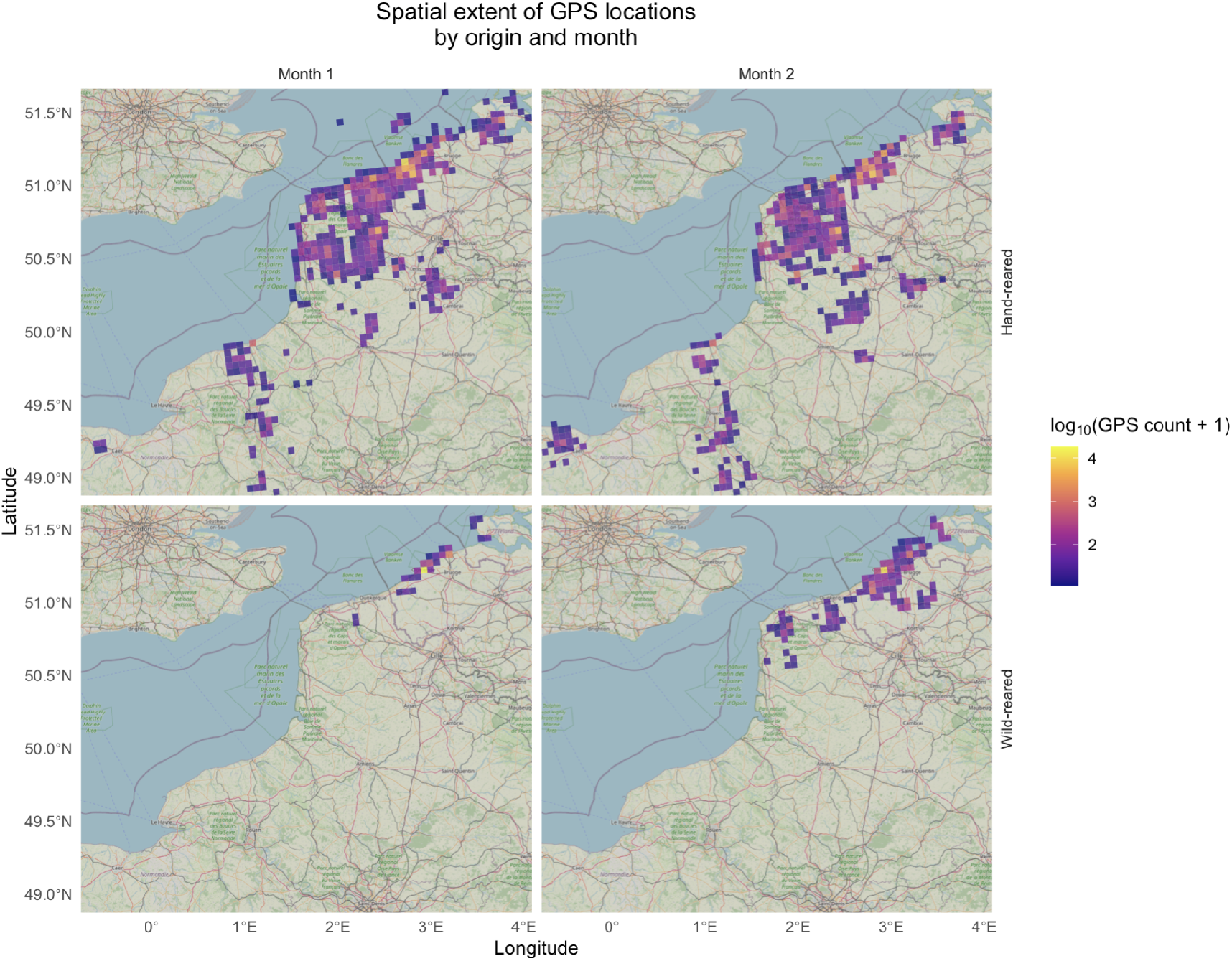
Spatial extent of GPS-positions over time for both groups.

To test for differences in occurrence distribution (i.e. space use) between hand- and wild-reared juveniles over time, we fitted a generalised least squares (GLS) model using the `gls()` function in `nlme` (Pinheiro and DM Bates, 2000). The model included group (hand-vs. wild-reared), window index, and their interaction as fixed effects. An AR(1) correlation structure accounted for temporal autocorrelation among successive windows within each bird. Residual plots and variogram checks confirmed that the AR(1) structure adequately captured within-individual dependence and that residuals met assumptions of normality and homoscedasticity.

#### Prediction 3: Occurrence near humans

To quantify each bird’s association with human population density, we extracted population data from the Global Human Settlement (GHS) population grid for the year 2020 at a 100m spatial resolution (**jrc_ghs**). We averaged population-density values across all daytime fixes within each of the two 30-day blocks after fledging. Cells with missing population-density values (e.g. sea or otherwise uninhabitable areas) were set to zero prior to averaging. These per-bird, per-month mean densities were analysed in a linear mixed-effects model with fixed effects of *group* (hand-reared vs. wild-reared), *month block* (first vs. second 30-day block) and their interaction, and a random intercept for individual, using `lmer()` in the `lme4` package (D Bates et al., 2015). A log(1 + *x*) transformation of the response was used to improve residual structure.

#### Prediction 4a: Habitat use

For habitat-use analyses we used the same daytime-only GPS fixes, excluding all positions within the origin-site area. Raw CORINE land-cover codes were recoded into three biologically meaningful categories: *Rural green areas, Marine areas*, and an *Anthropogenic areas* category that pooled all remaining (primarily human-modified) classes (see Supplementary Table 1). While the *Marine* category was defined to cover all water-related habitats, the majority of fixes within this group occurred specifically within the intertidal zone. Similarly, we pooled all human-modified classes into a single *Anthropogenic* category because individual classes were relatively rare in our study area, this aggregation avoided sparsely populated levels while preserving a clear contrast between semi-natural and human-modified habitats.

For each bird and each 30-day block, we tallied counts of daytime fixes in each habitat category in a given month. We then fitted a count GLMM for allocation among habitats using `glmmTMB` (McGillycuddy et al., 2025). The model used a generalised Poisson family (log link) with fixed effects of *group, habitat category, month block*, and all interactions, a log-offset for each bird’s total non-origin fixes within the corresponding 30-day block, and a random intercept for individual.

#### Prediction 4b: Habitat selection

We used the same daytime-only GPS tracks to perform integrated step-selection analyses (iSSA) via the `amt` package (Signer et al., 2019), using the same preprocessing steps as in the ‘Habitat use’ analysis. Due to data constraints, we had to collapse the data across all 60 days for this analysis.

For each individual, observed steps were defined between consecutive 30-minute fixes, and for every observed step, 30 random steps were generated using `random_steps()`, with step lengths and turning angles drawn from fitted gamma and von Mises distributions, respectively (Avgar et al., 2016). We then fitted an iSSF including habitat and standard movement covariates, and used `log_rss()` to obtain log-relative selection strengths (log-RSS) for each focal habitat (Rural green areas, Marine areas) relative to Anthropogenic. Log-RSS values were evaluated at the population median step length and mean turning angle. We modelled these individual log-RSS estimates in a weighted linear mixed-effects model fitted with `lmer()` in lme4 (D Bates et al., 2015), with *p*-values obtained from `lmerTest` (Kuznetsova et al., 2017). The fixed-effects structure included habitat (Rural green vs. Marine), group (hand-reared vs. wild-reared) and their interaction, with a random intercept for individual and inverse-variance weights. Weights were proportional to the inverse squared standard error of each log-RSS estimate (scaled to have mean 1), so that more precise estimates contributed more strongly to the fixed effects (see Supplementary Materials for full model specification and weighting details).

Habitat selection was modelled as

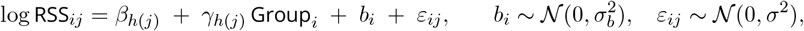

fitted by REML with precision weights *w*_*ij*_.

Here, *Groupi* is the group indicator for individual *i, h*(*j*) denotes the habitat associated with observation *j*, the coefficients *β*_*h*_ represent the expected log-RSS for habitat *h* in hand-reared juveniles (selection relative to *Anthropogenic*), and *γ*_*h*_ represent the within-habitat difference between wild-reared and hand-reared birds on the log scale. The random intercept *b*_*i*_ accounts for individual-level variation, and *w*_*ij*_ are inverse-variance weights (proportional to 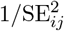, scaled to mean 1).

### Survival (Prediction 5)

We assessed two-month post-fledging survival using binomial logistic regression fitted with `glm()` in the `stats` package. Continuous GPS tracking provided unambiguous status at day 60, so no right-censored time-to-event models were needed. To verify that early deaths, potentially caused by carryover effects of GPS-tagging, did not bias our results we ran two analyses: one excluding and one including individuals that died within three days of fledging. Birds with tag failures during the 60 days and those classified as *potentially dead* were excluded (see Table 1, for classification criteria). Survival status was modelled as a binomial response with a logit link, including *Group* (hand-reared vs. wild-reared) as the main predictor. *Body_Condition* (mean-centered residuals of mass on tarsus length at fledging; Rotics et al. 2021) and its interaction with *Group* were added as covariates to control for potential effects of condition at fledging.

## 3 Results

### Behaviour (Predictions 1–4)

#### Prediction 1: Proportion of daytime fixes spent at site of origin

Using a binomial GLMM (*n* = 87 birds; 174 bird–months), we found strong effects of group and time on the proportion of daytime fixes within 500 m of the origin site. Relative to hand-reared birds, wild-reared juveniles had much higher odds of returning to their site of origin (*β*_wild-reared_ = 4.85 ± 0.34, *z* = 14.18, *p* < 0.001; OR ≈ 128 for wild-reared vs. hand-reared). Both groups reduced visits to the site of origin in the second month (*β*_month2_ = −2.14 ± 0.02, *z* = −88.78, *p* < 0.001; OR ≈ 0.12). Back-transformed model estimates indicate that in month 1, hand-reared juveniles spent ~2.4% of daytime fixes at the site of origin versus ~76.0% for wild-reared juveniles, declining in month 2 to ~0.3% (hand-reared) versus ~27.0% (wild-reared; Figure 3b).

**Figure 3.**
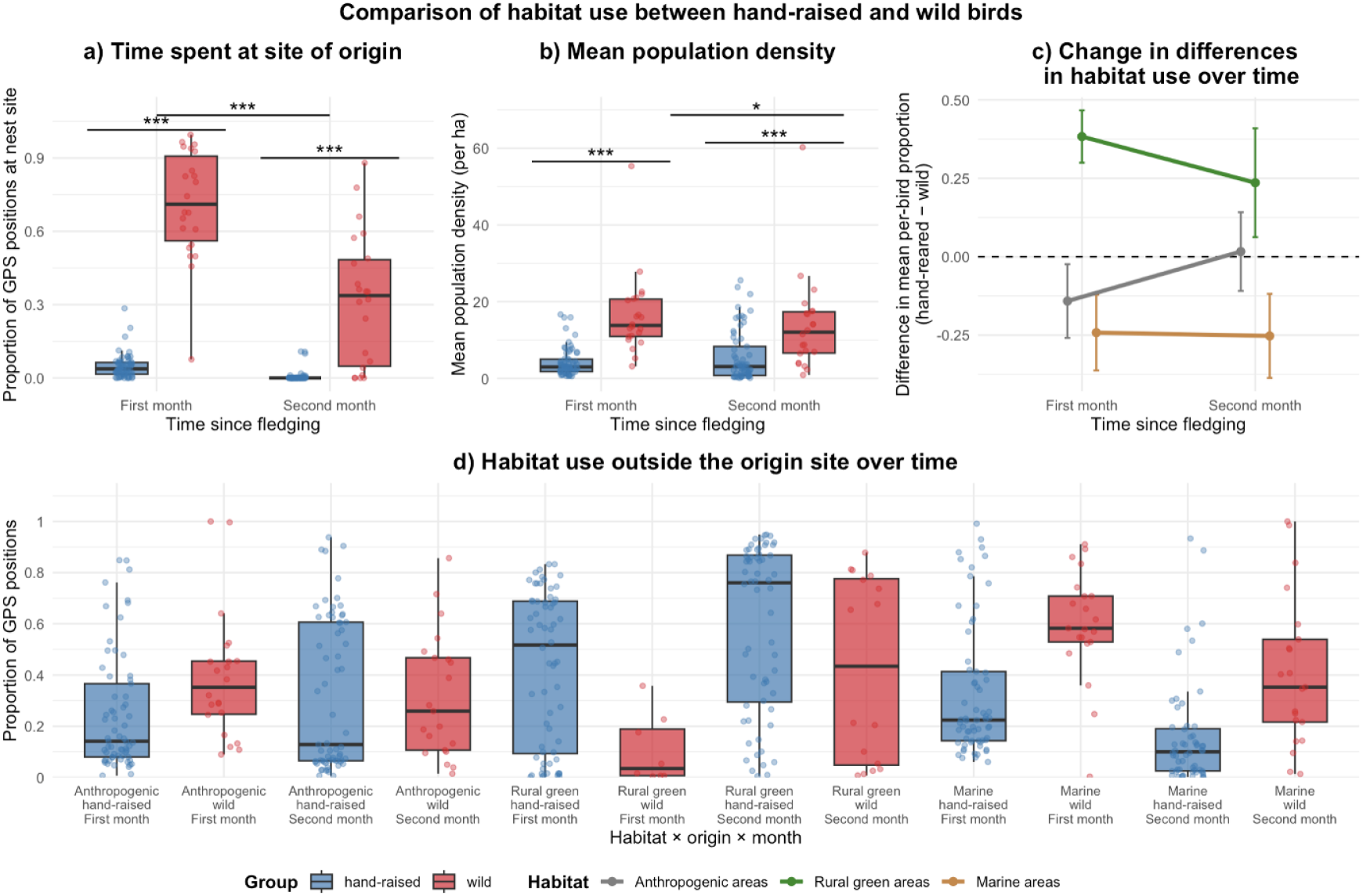
Differences in behaviour between hand-reared and wild-reared juveniles during the first 60 days after fledging, shown separately for the first and second 30-day blocks (“first” and “second” month). **(a)** Mean human population density (people/ha) at each bird’s daytime GPS locations. **(b)** Time spent at the origin site: proportion of daytime GPS fixes within 500 m of the origin site (release site for hand-reared, nest for wild-reared). **(c)** Change in between-origin differences in habitat use outside the origin site: for each habitat category, lines show the difference in mean per-bird proportion of non-origin fixes (hand-reared minus wild-reared); positive values indicate greater use by hand-reared birds. **(d)** Habitat use outside the origin site: per-bird proportion of daytime non-origin fixes in three lumped habitat categories (Anthropogenic areas, Rural green areas, Marine areas) by origin and month. Horizontal brackets in panels (a–c) indicate significant contrasts between groups or months (*** *p* < 0.001, ** *p* < 0.01, * *p* < 0.05; Holm-adjusted where applicable). For panel (d), significance labels were omitted.

#### Prediction 2: Space use

A GLS model of log-transformed 95% home range area across 51 rolling 10-day windows per bird (n = 87) revealed a strong group difference in initial space use: hand-reared juveniles started with larger occurrence distributions than wild-reared birds (*β* = −5.95 ± 0.58, *t* = −10.17, *p* < 0.001; Supplementary figure 4). Over time, hand-reared birds showed a gradual decline in occurrence distribution (*β* = −0.02 ± 0.01, *t* = −3.34, *p* < 0.001), whereas wild-reared juveniles expanded their space use (*β* = 0.10 ± 0.01, *t* = 7.31, *p* < 0.001).

**Figure 4.**
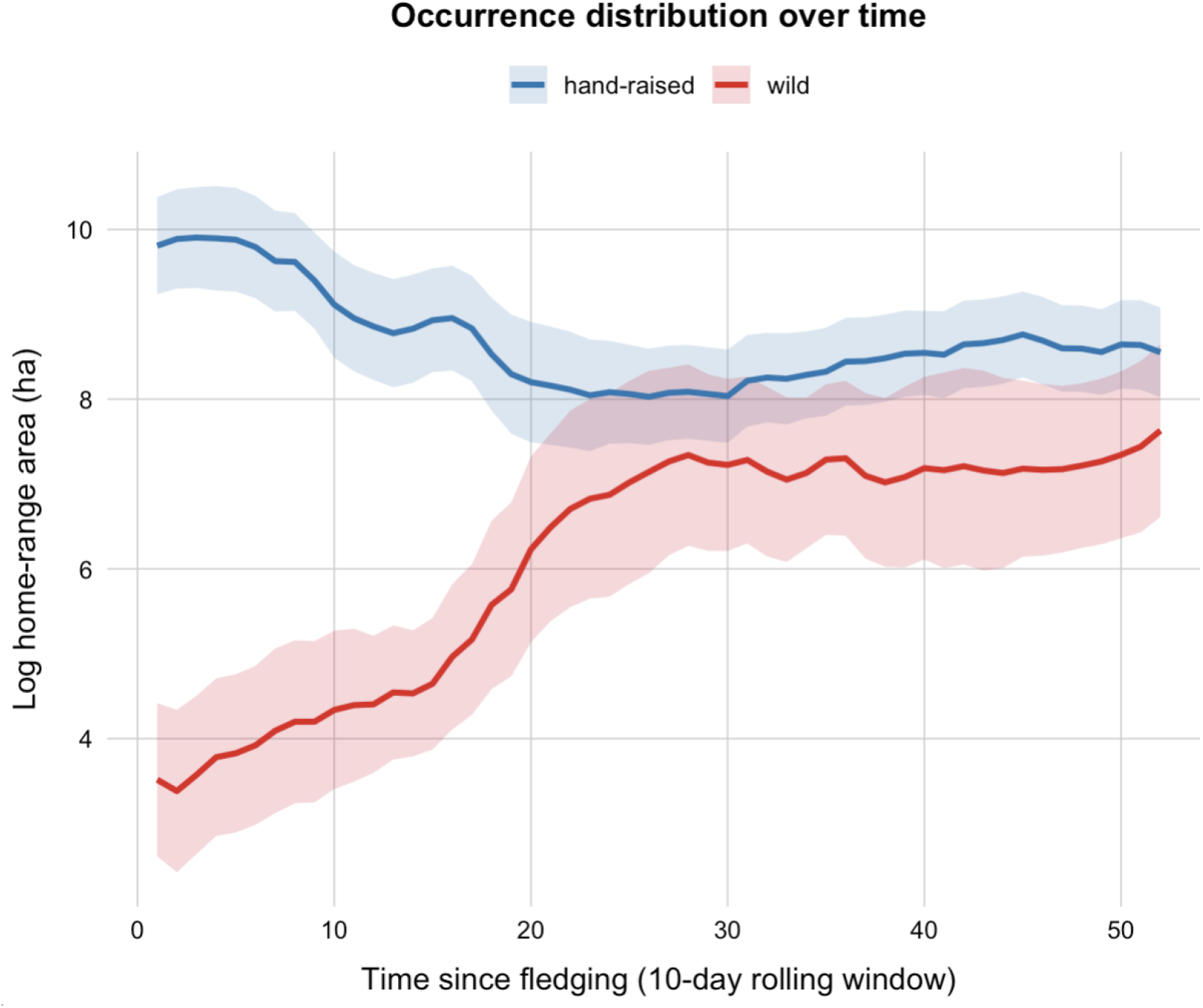
Change in log-transformed occurrence distribution area (95% MCP) over time in hand-reared and wild-reared juvenile herring gulls. Values represent mean area per 10-day window with 95% confidence intervals.

#### Prediction 3: Occurrence near humans

Using a linear mixed-effects model on log-transformed mean human population density, we found a strong effect of group and a group–by–month interaction (Figure 3a). Relative to hand-reared juveniles, wild-reared birds occurred in substantially more densely populated areas (*β*_wild-reared_ = 1.26 ± 0.19, *t* = 6.70, *p* < 0.001). There was no overall main effect of month (*β*_month2_ = −0.01 ± 0.08, *t* = −0.08, *p* = 0.94), but the significant interaction term indicated that the group difference was smaller in the second month (*β*_wild-reared×month2_ = −0.30 ± 0.15, *t* = −2.00, *p* = 0.048).

Back-transformed model estimates showed that in the first 30-day block, hand-reared juveniles used locations with an average of ~3.3 people/ha (95% CI: 2.6–4.2), compared with ~14.2 people/ha (10.0–19.9) for wild-reared juveniles. In the second block, hand-reared juveniles again averaged ~3.3 people/ha (2.6–4.2), whereas wild-reared juveniles shifted to somewhat less densely populated areas, averaging ~10.1 people/ha (7.1–14.3).

#### Prediction 4a: Habitat use

Post hoc contrasts from the habitat-use model (Supplementary Table 3) revealed group differences in the first 30 days after fledging (Figure 3c). In month 1, hand-reared juveniles allocated a much larger share of their time to rural green areas than wild-reared birds (hand-reared/wild-reared rate ratio ≈ 6.8, *z* = 5.00, *p* < 0.001), whereas wild-reared juveniles concentrated more strongly on anthropogenic and marine habitats (hand-reared/wild rate ratios ≈ 0.39 and ≈ 0.42, respectively; anthropogenic: *z* = −4.61, *p* < 0.001; marine: *z* = −3.97, *p* < 0.001). In month 2, hand-reared juveniles still used rural green areas more often than wild-reared birds (hand-reared/wild rate ratio ≈ 3.4, *z* = 4.30, *p* < 0.001), and wild-reared juveniles continued to use marine habitats more (ratio ≈ 0.27, *z* = −5.25, *p* < 0.001), whereas the origin difference in anthropogenic areas had weakened and was no longer supported statistically (*z* = −1.33, *p* = 0.184). Thus, while habitat use converged to some extent over time, wild-reared juveniles remained more strongly associated with anthropogenic and marine habitats than hand-reared birds (Figure 3d).

#### Prediction 4b: Habitat selection

Post hoc contrasts from the habitat-selection model, expressed as log-RSS relative to anthropogenic areas, revealed clear differences between groups in how strongly they selected habitats over a 60-day period. Relative to anthropogenic habitats, hand-reared juveniles avoided rural green areas (RSS ≈ 0.49; 95% CI: 0.39–0.61), whereas wild-reared juveniles showed no clear preference for rural green areas (RSS ≈ 1.33; 95% CI: 0.84–2.12, CI spans 1). This between-group contrast was significant (hand-reared − wild-reared estimate = −1.01 ± 0.26, *t*_69.2_ = −3.86, *p* < 0.001). For marine areas, hand-reared juveniles showed no clear preference relative to anthropogenic habitats (RSS ≈ 1.07; 95% CI: 0.87–1.32), whereas wild-reared juveniles clearly selected marine areas (RSS ≈ 1.75; 95% CI: 1.18–2.59). The group contrast was again significant (hand-reared − wild-reared estimate = −0.49 ± 0.22, *t*_92.2_ = −2.19, *p* = 0.031).

Although hand-reared juveniles were frequently located in rural green areas, the step-selection model indicated that, given what was locally available, they chose anthropogenic areas even more often than rural green areas. Wild-reared juveniles showed no clear difference between rural green and anthropogenic habitats, but selected marine habitats over anthropogenic ones.

### Survival (Prediction 5)

Two-month post-fledging survival did not differ significantly between hand- and wild-reared juveniles, nor was it predicted by body condition at the time of GPS-tagging. When early mortalities (juveniles that died within three days of fledging) were included (n = 198; 144 hand-reared, 54 wild-reared), 66 of 144 hand-raised juveniles (45.8%) and 37 of 54 wild-reared juveniles (68.5%) survived to day 60. Logistic regression revealed no significant effect of group (hand-raised vs. wild-reared:*β* = 0.37 ± 0.53, *z* = 0.70, *p* = 0.480; OR = 1.45, 95% CI: 0.51–4.14), body condition at fledging (*β* = 0.001 ± 0.002, *z* = 0.43, *p* = 0.670), or their interaction (*β* = −0.001 ± 0.006, *z* = −0.13, *p* = 0.900).

Excluding juveniles that died within three days of fledging (n = 184; 130 hand-raised, 54 wild-reared) produced similar results: 66 of 130 hand-raised (50.8%) and 37 of 54 wild-reared (68.5%) survived to day 60. Again, rearing origin was not a significant predictor (*β* = 0.11 ± 0.54, *z* = 0.20, *p* = 0.840; OR = 1.12, 95% CI: 0.39–3.17), nor was body condition (*β* = 0.000 ± 0.002, *z* = 0.20, *p* = 0.840) or their interaction (*β* = −0.001 ± 0.006, *z* = −0.10, *p* = 0.920). Full model outputs, including all coefficient estimates, odds ratios, and confidence intervals, are provided in Supplementary Table 2.

## 4 Discussion

To understand how parental attendance after fledging relates to early movement, habitat use and selection, and survival, we compared hand-reared juveniles with wild-reared birds. This contrast necessarily also includes differences in parental care during the nestling phase, and we acknowledge that such early-life effects could contribute to later behaviour. However, because our analyses begin at fledging and focus on the establishment of independent space use and foraging, we interpret patterns primarily in relation to post-fledging attendance, guidance, and provisioning by parents unless stated otherwise. Hand-reared juveniles left their origin site sooner, ranged more widely, and occurred in areas with lower human population density. Their habitat use was dominated by rural green habitats, whereas wild-reared birds used anthropogenic and marine habitats more strongly early on. At the same time, the habitat-selection analysis indicated that hand-reared juveniles avoided rural green landscapes relative to availability, suggesting that their broader space use did not translate into proportional selection for farmland at fine scales. Although spatial behaviour and habitat use of the two groups tended to converge over the first two months, distinct differences remained. Yet, despite these contrasts in space use and habitat associations, we detected no difference in short-term survival between the groups.

Consistent with Predictions 1–2, hand-reared juveniles spent less time at their site of origin and initially ranged over larger areas than wild-reared birds (Figure 2,4). In the first month after fledging, wild juveniles still concentrated most of their daytime activity near the natal site, whereas hand-reared birds, released without attending parents, spent less time at their release site and distributed their movements more widely. By the second month, time at the origin site had decreased sharply in both groups. Over the same period, home-range estimates showed that wild-reared juveniles expanded their space use, consistent with increasing independence and reduced parental attendance as juveniles separate from their parents, whereas hand-reared birds contracted their ranges, consistent with settlement after an initial broad search phase, so that range sizes converged. The initial broad ranging and subsequent contraction observed in hand-reared juveniles is a pattern also reported in reintroduction programmes where animals are released into unfamiliar terrain (Kemp et al., 2020; Ram et al., 2025; Reyna et al., 2021). In contrast, wild-reared juveniles showed the opposite trend, gradual expansion, likely because initial post-fledging parental care dampens early exploratory movements. By providing food near the nest and guiding movements (Holley, 1982; Spear et al., 1986), parents can anchor wild juveniles to the wider natal area, delaying their transition to broader space use. Since our analyses excluded movements within 500 m of the origin, these diverging patterns likely reflect behavioural consequences of (the absence of) parental care rather than simple differences in flight capability.

Contrary to Prediction 3, hand-reared juveniles were less likely to be found in areas with high human population density compared with wild-reared birds, particularly during the first month postfledging. Although wild-reared juveniles also moved to slightly less densely populated areas in the second month, the difference between the groups persisted. This pattern matches colour-ring resighting data from the same system, where wild juveniles were observed in areas of intense human activity more frequently than rehabilitated ones (Uytterschaut et al., 2025). In line with Prediction 4a, habitat use also differed clearly between origins. Hand-reared juveniles primarily used agricultural habitats, whereas wild-reared juveniles favoured anthropogenic and marine areas. Over time, these profiles converged somewhat: both groups increased their use of agricultural areas and decreased their use of marine zones. Additionally, wild-reared birds reduced their reliance on anthropogenic areas, narrowing, but not eliminating, the difference between the two groups.

These patterns suggest that post-fledging parental care may buffer early constraints, allowing wild-reared juveniles to remain in profitable but highly competitive environments, even though other processes may also contribute. Foraging in cities or restricted marine zones (which along the Belgian coast are limited to narrow, crowded breakwaters in the intertidal zone) involves intense competition and a more complex risk landscape for inexperienced juveniles (Donk et al., 2019; Enners et al., 2018). Wild-reared juveniles may persist in these settings for longer if parental attendance and provisioning reduce the costs of low foraging competence and promote learning in familiar areas, although we did not quantify parental provisioning directly. In contrast, hand-reared juveniles are forced to become independent immediately and may therefore shift sooner towards agricultural landscapes, where resources are more diffuse and profitable opportunities can often be located by tracking conspicuous activity, such as aggregations of gulls around farming operations. The later increase in farmland use by wild-reared birds may reflect growing independence, but it could also be partly seasonal, as post-fledging periods extend into late summer and autumn when agricultural disturbance and resource availability change (for instance during harvesting, ploughing, or manure spreading), creating short-lived foraging opportunities.

The association of wild-reared juveniles with higher human population densities may partly be a byproduct of overall habitat use, but it could also reflect nest-origin and coastal geography. Ostend is a seaside town where beaches, harbour infrastructure and shoreline features lie within, or directly adjacent to, anthropogenic habitat, so coastal habitat use can coincide with high population densities even without a direct attraction to humans. Hand-reared individuals, by contrast, dispersed early into the rural agricultural landscapes of northern France (Figure 2) where human density is lower, so the population-density contrast likely reflects their specific habitat use rather than a direct preference for humans.

Although our habitat-use analysis showed that hand-reared juveniles spent the majority of their time in agricultural landscapes, our habitat-selection model reveals that they still statistically avoided this habitat relative to anthropogenic areas. This suggests that while hand-reared birds are often found in agricultural environments, they disproportionately utilise anthropogenic features in this landscape, such as waste management sites and industrial sites (with low human densities; see above), consistent with the well-documented use of landfill and other built-up habitats by herring gulls (Belant et al., 1993; O’Hanlon et al., 2022). Such sites may also be discovered quickly through conspecific attraction and local enhancement, because they generate strong, local aggregations of gulls (Thiebault et al., 2014; Weimerskirch et al., 2010). Indeed, exploratory visual analysis of individual GPS tracks revealed that several individuals spent a lot of time at a waste management site in northern France. While this site attracts a lot of adult gulls, the spatial layout may reduce direct interference competition if food is dispersed widely across the site, but we did not quantify competitive interactions.

It is important to note that our selection analysis pooled data over the first 60 days, providing an aggregate measure of preference that masks the temporal dynamics observed in the habitat use analysis. Consequently, while the habitat-use analysis revealed that both groups shifted their habitat use over the course of the study, leading to a convergence in their spatial patterns, the static selection model was unable to capture these dynamic trajectories.

Contrary to Prediction 5, we found no statistically significant difference in survival rates between the two groups. Overall survival fell within the typical first-year range reported for large gulls (~30–70% (Camphuysen and Gronert, 2012; Kentie et al., 2022; Schekkerman et al., 2021) and aligned with local estimates based on colour-ring resightings (Uytterschaut et al., 2025). However, although survival at day 60 was ≈18% lower in hand-reared (≈51%) than in wild-reared juveniles (≈69%), this contrast did not reach statistical significance. Given the wide confidence intervals, this result should be interpreted with caution, as it may reflect limited statistical power. It is also possible that costs related to a lack of parental care accrue gradually, becoming detectable only in autumn or winter as food availability declines and environmental conditions deteriorate (Cox et al., 2014). Finally, early postfledging mortality may have been driven by stochastic external hazards, such as the avian influenza outbreaks present during the study years (Lean et al., 2024), which could have masked subtler differences between the groups.

The finding that the absence of parental care profoundly shapes juvenile space use, even if shortterm survival remains comparable, is particularly relevant for rehabilitation and reintroduction programmes. In our data, these behavioural differences were most pronounced early after fledging and diminished over time, and this convergence is unlikely to be explained by selective disappearance because survival did not differ detectably between groups during the same period. It suggests that current release protocols may produce individuals that, while viable, are initially ecologically distinct from their wild counterparts. To bridge this behavioural gap, such programmes could implement staged provisioning to mimic the extended parental care that naturally anchors wild juveniles to high-quality habitat (e.g. soft release protocols). Additionally, fostering opportunities for social information use (e.g. housing with adults or targeted release near active colonies) might help hand-reared juveniles navigate the ‘exploration-exploitation’ trade-off more effectively, reducing their need to retreat into less competitive or suitable landscapes.

In interpreting our findings, some contextual and methodological limitations should be considered. Because hand-reared juveniles developed in captivity, some differences could in principle reflect captive-rearing experience rather than parental care alone. We reduced this as far as possible by applying comparable selection for tagging in both groups, focusing on individuals judged to be in the best condition, and by preparing hand-reared birds for release in ways intended to minimise association with humans and reduce predictable provisioning. Relatedly, our comparison is not a manipulation of a single component of parental care. Hand-reared juveniles lacked both pre- and post-fledging care, so we cannot separate these effects. Both can influence behaviour after fledging (Avery, 1996; Jones et al., 2025; Penttinen et al., 2025), but because our response variables capture the first weeks of independent movement, we focus on post-fledging attendance and provisioning as the most direct pathway shaping origin-site use, ranging behaviour and early habitat association. While our group-level models revealed clear differences in movement and habitat use, these population averages summarise considerable individual specialisation. Thus, our results describe central tendencies across diverse behavioural strategies rather than a single typical trajectory. Methodologically, a key constraint lies in the spatial context. Hand-reared juveniles were released in a single reserve, while wild-reared individuals originated from environmentally distinct sites (urban rooftops and ground-nesting colonies). Consequently, the observed habitat-use contrasts may reflect (especially in the first month) differences in local landscapes as much as differences stemming from parental care, suggesting a partial confounding of origin and spatial context.

Overall, by comparing hand-reared and wild-reared juveniles, we show that the absence of parental care markedly alters the pace and direction of early space use and habitat association, even though short-term survival does not differ detectably between groups. Longer-term monitoring is essential to determine whether these early behavioural differences persist, converge, or ultimately carry over to affect long-term survival, recruitment success, and overall fitness, because persistent shifts in settlement and habitat associations could have practical consequences for release design and post-release support in hand-rearing and reintroduction programmes.

## Author Contributions (CREDIT)

Conceptualization: R.A., L.L., F.V.; Methodology: R.A.; Data curation: R.A.; Formal analysis: R.A.; Visualization: R.A.; Validation: R.A.; Writing – original draft: R.A.; Writing – review and editing: R.A., J.V., W.M., E.S., A.M., L.L., F.V.; Supervision: L.L., F.V.; Funding acquisition: A.M., L.L., F.V.; Project administration: F.V.

## Acknowledgements

We are grateful to Hans Matheve (Ghent University) for his assistance with bird tagging. We thank Isabelle Allemeersch (Wildlife Rescue Centre Ostend) for valuable advice on hand-rearing, and the Gull Patrol team (Marc Verborgh and Nathan Noels) for providing eggs. Special thanks go to Nathan Audenaert for his invaluable help in raising the birds. Finally, we thank the Agentschap voor Natuur en Bos (ANB) for granting permission to release the birds at the IJzermonding nature reserve.

## Data, scripts, code, and supplementary information availability

All data required to replicate this study’s findings will be made openly available on Zenodo: https://doi.org/10.5281, code and raw data are accessible through the article’s OSF repository: https://osf.io/zpy3q/ Supplementary information supporting the results is also provided in this repository.

---

**Supplementary Table 1.** CORINE recoding.

| Recoded habitat category | Original CORINE class(es) |
| --- | --- |
| Anthropogenic areas | Continuous urban fabric; Discontinuous urban fabric; Airports; Construction sites; Industrial or commercial units; Road and rail networks and associated land; Port areas; Dump sites; Mineral extraction sites; Green urban areas; Sport and leisure facilities |
| Rural green areas | Broad-leaved forest; Coniferous forest; Mixed forest; Natural grasslands; Moors and heathland; Non-irrigated arable land; Pastures; Complex cultivation patterns; Land principally occupied by agriculture with significant areas of natural vegetation |
| Marine areas | Beaches; Dunes; Sands; Estuaries; Inland marshes; Intertidal flats; Salines; Salt marshes; Sea and ocean; Water bodies; Water courses |
| Nest site | Any fix within 500 m of the nest or release site |

## 5 Details of habitat-selection analysis

We computed three movement covariates for each step: step length `(sl_)`, log step length `(log_sl_)`, and the cosine of the turning angle `(cos_ta_)`, which were included as predictors. A separate `iSSF` was fitted for each individual using the formula:

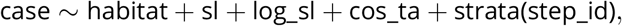

where `case_` indicates whether a step was observed (1) or random (0), `habitat` is a seven-level factor (see table 1, excluding “Nest site”), and `strata(step_id)` matches each observed step to its 30 random alternatives. From each fitted iSSF, we used `log_rss()` to extract log-relative selection strengths (log-RSS) for each habitat class, using “Rural green areas” as the reference. Log-RSS was evaluated at the median step length (131.25 m) and the mean cosine of the turning angle (0.107), computed across all steps. Any habitat–bird combination with a collapsed 95% CI was discarded, as this indicates zero encounters with that habitat.

**Supplementary Table 2.** Logistic regression coefficients, odds ratios (OR), 95% confidence intervals (CI), and p-values for filtered (excluding early deaths) and unfiltered (including early deaths) models predicting survival to 60 days post-fledging.

| Term | No early deaths (n = 132) |  |  | Early deaths (n = 146) |  |  |
| --- | --- | --- | --- | --- | --- | --- |
|  | Estimate (SE) | OR (95% CI) | p-value | Estimate (SE) | OR (95% CI) | p-value |
| Intercept | 0.125 (0.196) | 1.130 [0.770, 1.670] | 0.522 | −0.126 (0.184) | 0.880 [0.580, 1.200] | 0.494 |
| Group (wild) | 0.109 (0.541) | 1.110 [0.380, 3.360] | 0.841 | 0.372 (0.530) | 1.450 [0.510, 4.150] | 0.483 |
| Body_index | 0.001 (0.002) | 1.000 [0.996, 1.000] | 0.839 | 0.001 (0.002) | 1.001 [0.997, 1.003] | 0.668 |
| Group × body_index | −0.001 (0.006) | 0.999 [0.988, 1.010] | 0.922 | −0.001 (0.006) | 0.999 [0.988, 1.010] | 0.895 |
| Null Deviance | 182.23 on 131 df |  |  | 202.29 on 145 df |  |  |
| Residual Deviance | 182.13 on 128 df |  |  | 201.45 on 142 df |  |  |
| AIC | 190.13 |  |  | 209.45 |  |  |

For each remaining log-RSS estimate, standard errors were approximated as:

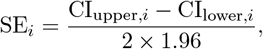

and inverse standard error weights were computed as:

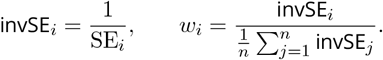

These weights were applied to give higher influence to habitat-selection estimates with greater precision (i.e., smaller SE), ensuring that individuals with more precise log-RSS estimates contribute more strongly to the fixed-effects estimates (Beardsworth et al., 2021; Graf et al., 2024).

**Supplementary Table 3.** GLMM and LMM results for time spent at the origin site (binomial), non-origin habitat allocation (generalised Poisson), and human population density (linear mixed-effects model).

| **Time at origin site:** $n\_fixes \sim status + month\_block + (1 \vert individual)$ (binomial GLMM, logit link) | | | | |
| --- | --- | --- | --- | --- |
| Term | Estimate | Std. Error | z value | Pr(> \|z\|) |
| (Intercept) | −3.704 | 0.176 | −21.05 | < 0.001 |
| Status<sub>wild</sub> | 4.852 | 0.342 | 14.18 | < 0.001 |
| Month<sub>2</sub> | −2.136 | 0.024 | −88.78 | < 0.001 |
| **Random-effects variance (Intercept):** 1.889 (Std. Dev. 1.375) |  |  |  |  |
| **Habitat use:** $n\_fixes \sim status * habitat\_3cat * month\_block + offset + (1 \vert individual)$ (generalised Poisson GLMM, log link) | | | | |
| Term | Estimate | Std. Error | z value | Pr(> \|z\|) |
| (Intercept) | −1.151 | 0.122 | −9.44 | < 0.001 |
| Status<sub>wild</sub> | 0.938 | 0.204 | 4.61 | < 0.001 |
| Rural green areas | −0.030 | 0.151 | −0.20 | 0.841 |
| Marine areas | 0.263 | 0.122 | 2.14 | 0.032 |
| Month<sub>2</sub> | −0.128 | 0.125 | −1.03 | 0.305 |
| Status<sub>wild</sub> × Rural green areas | −2.848 | 0.409 | −6.96 | < 0.001 |
| Status<sub>wild</sub> × Marine areas | −0.065 | 0.267 | −0.24 | 0.807 |
| Status<sub>wild</sub> × Month<sub>2</sub> | −0.647 | 0.266 | −2.43 | 0.015 |
| Rural green areas × Month<sub>2</sub> | 0.417 | 0.187 | 2.24 | 0.025 |
| Marine areas × Month<sub>2</sub> | −1.404 | 0.194 | −7.23 | < 0.001 |
| Status<sub>wild</sub> × Rural green areas × Month<sub>2</sub> | 1.330 | 0.519 | 2.56 | 0.010 |
| Status<sub>wild</sub> × Marine areas × Month<sub>2</sub> | 1.088 | 0.396 | 2.75 | 0.006 |
| **Reference habitat:** Anthropogenic areas. |  |  |  |  |
| **Random-effects variance (Intercept):** 0.144 (Std. Dev. 0.380) |  |  |  |  |
| **Dispersion parameter:** 576 |  |  |  |  |
| **Human population density:** $\log(1 + mean\_pop\_density) \sim status * month\_block + (1 \vert individual)$ (LMM) | | | | |
| Term | Estimate | Std. Error | t value | Pr(> \|t\|) |
| (Intercept) | 1.464 | 0.094 | 15.53 | < 0.001 |
| Status<sub>wild</sub> | 1.256 | 0.187 | 6.70 | < 0.001 |
| Month<sub>2</sub> | −0.006 | 0.077 | −0.08 | 0.941 |
| Status<sub>wild</sub> × Month<sub>2</sub> | −0.305 | 0.152 | −2.00 | 0.048 |
| **Random-effects variance (Intercept):** 0.387 (Std. Dev. 0.622); Residual variance 0.190 (Std. Dev. 0.436) |  |  |  |  |
| **Back-transformed model-estimated means (people/ha) by origin and month block** |  |  |  |  |
| Group | Month block | Mean | 95% CI | n |
| Hand-reared | First 30 days | 3.3 | [2.6, 4.2] | 65 |
| Wild-reared | First 30 days | 14.2 | [10.0, 19.9] | 22 |
| Hand-reared | Days 31–60 | 3.3 | [2.6, 4.2] | 65 |
| Wild-reared | Days 31–60 | 10.1 | [7.1, 14.3] | 22 |

